# Modeling Sex Differences in the Effects of Diuretics in Renal Epithelial Transport during Angiotensin II-induced Hypertension

**DOI:** 10.1101/2023.12.11.571093

**Authors:** Kaixin Zheng, Anita T. Layton

## Abstract

Chronic angiotensin II (AngII) infusion is an experimental model that induces hypertension in rodents. The natriuresis, diuresis, and blood pressure responses differ between males and females, perhaps unexpectedly, given the rodent kidney, which plays a key role in blood pressure regulation, exhibit marked sex differences. Those sex differences include morphology, hemodynamics, and, under healthy (undrugged) conditions, solute and electrolyte transporter abundance. Notably, compared to the male rat nephron, the female rat nephron exhibits lower Na^+^/H^+^ exchanger 3 (NHE3) activity along the proximal tubule, but higher Na^+^ transporter activities along the distal segments. AngII infusion-induced hypertension induces a pressure natriuretic response that reduces NHE3 activity and shifts Na^+^ transport capacity downstream, to different extents in the two sexes. The goals of this study are (i) to understand how the sexually dimorphic responses differentially impact segmental electrolyte transport following a 14- day AngII infusion, and (ii) to identify and explain any sex differences in the effects of loop diuretics, thiazide diuretics, and K^+^-sparing diuretics. To achieve those goals, we developed sex-specific computational models of renal epithelial transport of electrolytes and water. Model simulations suggest that the NHE3 downregulation in the proximal tubule is a major contributor to natriuresis and diuresis in hypertension, with a stronger effect in males. Due to the downstream shift of Na^+^ transport load in hypertension, all three diuretic classes are predicted to induce stronger natriuretic and diuretic effects under hypertension compared to normotension, especially in females.

**New and Noteworthy:** Sex differences in the prevalence of hypertension are found in humans and animal models. The kidney, which plays an important role in blood pressure regulation, exhibits sex differences in morphology, hemodynamics, and membrane transporter distributions. This computational modeling study provides insights into how the sexually dimorphic responses to a 14-day angiotensin II infusion differentially impact segmental electrolyte transport. Simulations results also explain sex differences in the effects of loop diuretics, thiazide diuretics, and K^+^-sparing diuretics.

## Introduction

The renin-angiotensin-aldosterone system (RAAS) contributes to the regulation of blood volume, electrolyte balance, and systemic vascular resistance (1). An overactive RAAS may lead to hypertension. RAAS blockers, notably angiotensin-converting enzyme (ACE) inhibitors and angiotensin II (AngII) receptor blockers, are widely prescribed anti-hypertensive therapies. On the other hand, chronic AngII infusion is a well-studied model of experimental hypertension (2). A 14-day infusion of AngII induces vasoconstriction and anti-natriuresis, resulting in elevated blood pressure (3,4). At a molecular level, a 14-day infusion of ANGII in rats decreased the abundance of Na^+^/H^+^ exchanger 3 (NHE3) in the proximal tubule and that of Na^+^-K^+^-Cl^-^ cotransporter 2 (NKCC2) in the medullary thick ascending limb. Interestingly, the abundance of NKCC2 is increased in the cortical thick ascending limb, as is the abundance of Na^+^- Cl^−^ cotransporter (NCC) in the distal convoluted tubule, and epithelial Na^+^ channel (ENaC), which is expressed in the convoluted tubule and throughout the collecting duct.

Given AngII’s multitude of effects on kidney function, substantial efforts have been invested to attain a better understanding of how AngII regulates renal Na^+^ transport at a molecular level, and how that regulation affects males and females differently. Veiras et al. (5) documented variations in transporter levels between male and female (normotensive) rodents. Their results highlighted significant differences in transport capacity within the proximal tubule of male and female rat nephrons.

Specifically, when compared to male rat nephrons, female rat nephrons exhibited increased NHE3 phosphorylation in the proximal tubule, along with a shift in distribution towards the base of the microvilli. Additionally, female rat nephrons showed reduced abundance of Na^+^-Pi cotransporter 2 (NaPi2), aquaporin 1 (AQP1), and claudin-2 compared to male rat nephrons. In contrast, beyond the macula densa, female rat nephrons displayed higher levels of both abundance and phosphorylation of NCC, as well as increased abundance of ENaC and claudin-7 compared to male rats. Given these sex differences in renal transporter expression pattern, how might AngII’s regulation of renal Na^+^ transport differ between the sexes?

Sex differences have been reported in rodents in their response to chronic administration of AngII. Xue et al. (6) observed a 5-fold larger increase in blood pressure in male than female mice following 7-day infusion of AngII. Toering et al. (7) measured the changes in blood pressure and proteinuria in response to chronic administration (200 ng/kg/min for 3 weeks) of AngII in rats. They found that male rats showed a more rapid increase in systolic blood pressure than female rats and had an increased response of proteinuria compared with females.

Diuretics have a longstanding history as a treatment of hypertension, often employed as a first-line therapeutic option in clinical practice and form an important component of multiple-combination therapies (8,9). Acting on the kidneys, diuretics impair the activity of Na^+^ transporters along specific segments of the nephron, leading to increased urinary water and solutes excretions and lowered blood pressure (10). In this study, the natriuretic and diuretic effects of three classes of diuretics on a kidney in AngII-induced hypertension are investigated: loop diuretics, thiazide diuretics, and K^+^- sparing diuretics. These three classes of diuretics target different Na^+^ transporters and segments: loop diuretics inhibit the Na^+^-K^+^-2Cl^-^ cotransporter 2 on the apical membrane of thick ascending limb; thiazide diuretics, Na^+^-Cl^-^ cotransporter along distal convoluted tubule; and K^+^-sparing diuretics, ENaC along the distal convoluted tubule and collecting duct. Thiazide diuretics are well-studied and popular, due to its safety and effectiveness in both experimental studies and clinical practice (11,12). K^+^-sparing diuretics, alone or combinedly applied with thiazide, are also one of the main diuretic treatments (13,14).

There are known sex differences in the development of AngII-induced hypertension in mice (6,15). Nonetheless, the precise mechanisms by which those sex differences arise from the sexually dimorphic renal transporter patterns are incompletely known. Thus, we seek to employ modeling analysis to investigate the following: How do sex differences in renal transporter patterns impact the kidney’s response to chronic AngII infusion? How do the natriuretic and diuretic effects of diuretics differ between the sexes? To answer these questions, we develop sex-specific computational models of epithelial transport in a nephron of a rat kidney under AngII-induced hypertension and simulate the effects of diuretics for both sexes.

## Methods

### Model structure

We simulate the effect on kidney function in male and female rats of hypertension induced by 14-day infusion of AngII. This is done by adapting published sex-specific computational models of solute and water transport through epithelial cell in the superficial nephron of the rat kidney (16,17). The model nephron is divided into several functionally distinct segments: proximal convoluted tubule (PCT), proximal straight tubule (S3), short descending limb (SDL), medullary thick ascending limb (mTAL), cortical thick ascending limb (cTAL), distal convoluted tubule (DCT), connecting tubule (CNT), cortical collecting duct (CCD), outer medullary collecting duct (OMCD), and inner medullary collecting duct (IMCD). Each segment is modelled with different physical dimensions, transporter profile, permeability, and features like coalescence. In each computational cell of individual segment, steady state luminal, cellular, and paracellular concentrations and fluxes are calculated based on water conservation, non-reacting solute conservation, and pH conservation. Model parameters can be specified to simulate nephron function in a male or female rat kidney, under normotensive or hypertensive conditions; see Table 1. The model includes transporters shown in schematic diagram Fig.1.

**Figure 1.**
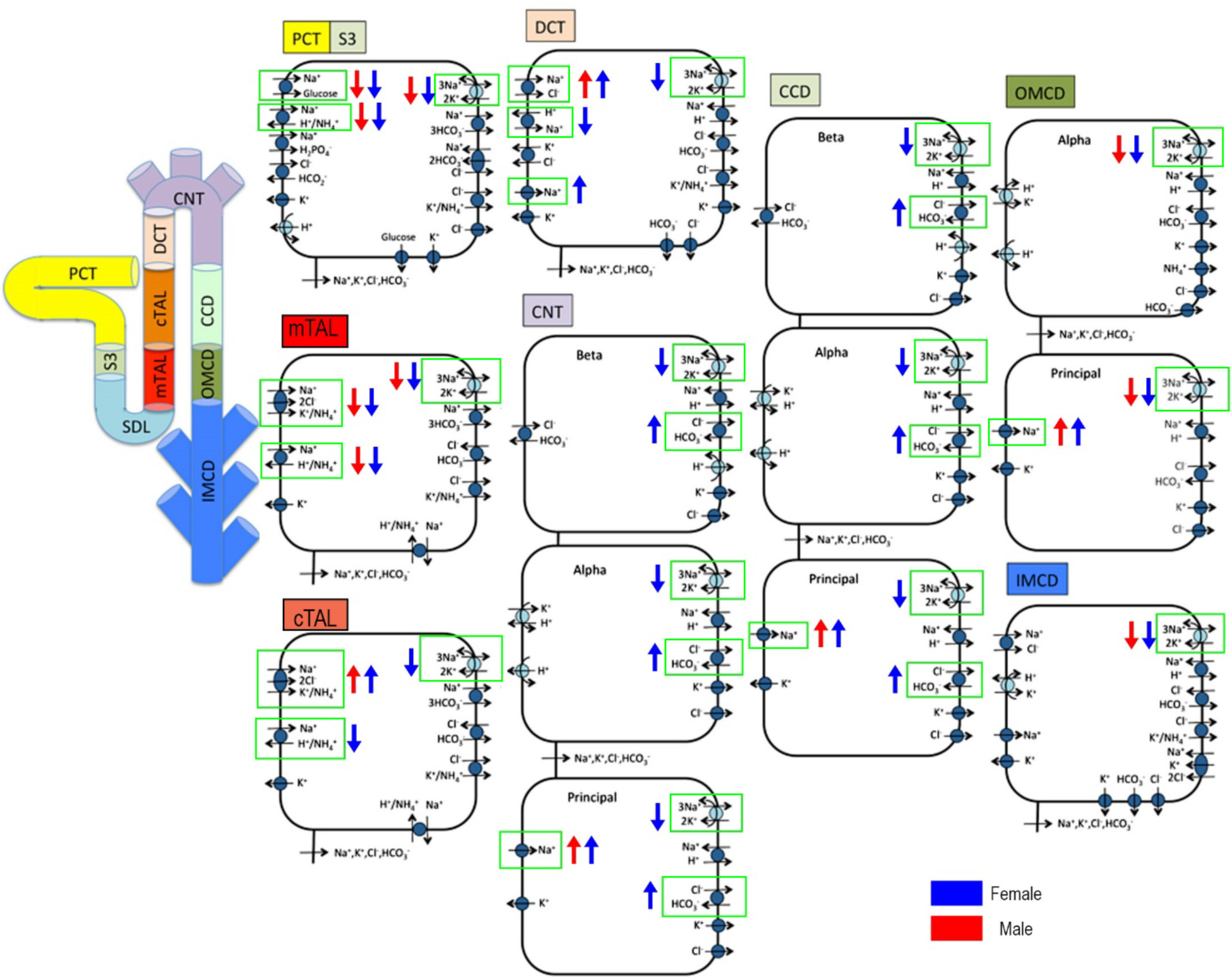
Schematic diagram of a superficial nephron in a rat kidney, with its epithelial cell components. The model accounts for the transport of 15 solutes and water. The cell diagrams show primarily Na^+^, K^+^, and Cl^-^ transporters. PCT, proximal convoluted tubule; S3, proximal straight tubule; SDL, short descending limb; mTAL, medullary thick ascending limb; cTAL, cortical thick ascending limb; DCT, distal convoluted tubule; CNT, connecting tubule; CCD, cortical collecting duct; OMCD, outer medullary collecting duct; IMCD, inner medullary collecting duct. Up/down arrow indicates transporter upregulation/downregulation due to hypertension. Adapted from Ref. (16).

**Table 1.** Hypertensive model parameter adaptations. HTN, hypertension; NTN, normotension; NHE3, Na^+^ /H^+^ exchanger isoform 3; ENaC, epithelial Na^+^ channel; NKCC2, Na^+^-K^+^-2Cl^-^ cotransporter 2 (A, B, F indicates isoform type); NCC, Na^+^-Cl^-^ cotransporter; SGLT1, sodium-glucose cotransporter 1; SGLT2, sodium-glucose cotransporter 2; Pf, water permeability; Purea, urea permeability.

|  | HTN-to-NTN ratios |  |
| --- | --- | --- |
| Parameter | Male | Female |
| **Proximal convoluted tubule** |  |  |
| NHE3 | 0.78 (3) | 0.79 (4) |
| Na<sup>+</sup>-K<sup>+</sup>-ATPase | 1 | 0.8 (22) |
| SGLT2 | 1 | 0.65 (20) |
| **S3** |  |  |
| NHE3 | 0.55 (3) | 0.67 (4) |
| Na<sup>+</sup>-K<sup>+</sup>-ATPase | 1 | 0.8 (22) |
| SGLT1 | 1 | 2.05 (20) |
| **Medullary thick ascending limb** |  |  |
| NHE3 | 0.55 (3) | 0.67 (4) |
| NKCC2A | 0.83 (3) | 0.70 (4) |
| NKCC2F | 0.83 (3) | 0.70 (4) |
| Na<sup>+</sup>-K<sup>+</sup>-ATPase | 0.7 (3) | 0.62 (4,22) |
| **Cortical thick ascending limb** |  |  |
| NKCC2A | 1.74 (3) | 1.50 (4) |
| NKCC2B | 1.74 (3) | 1.50 (4) |
| NHE3 | 1 | 0.88 (4) |
| Na<sup>+</sup>-K<sup>+</sup>-ATPase | 1 | 0.80 (22) |
| **Distal convoluted tubule** |  |  |
| NCC | 1.93 (3) | 2 (4) |
| Na<sup>+</sup>-K<sup>+</sup>-ATPase | 1 | 0.80 (22) |
| NHE3 | 1 | 0.88 (4) |
| ENaC | 1 | 2.42 (4) |
| **Connecting tubule** |  |  |
| ENaC | 1.21 (3,18) | 2.42 (4) |
| Na<sup>+</sup>-K<sup>+</sup>-ATPase | 1 | 0.8 (22) |
| apical P<sub>f</sub>, principal cell | 1 | 0.5 (20) |
| pendrin | 1 | 1.47 (20) |
| **Cortical collecting duct** |  |  |
| ENaC | 1.21 (3,18) | 2.42 (4) |
| Na<sup>+</sup>-K<sup>+</sup>-ATPase | 1 | 0.8 (22) |
| pendrin | 1 | 1.47 (20) |
| apical P<sub>f</sub>, principal cell | 1 | 0.5 (20) |
| **Outer-medullary collecting duct** |  |  |
| ENaC | 1.21 (3,18) | 1.57 (4,20) |
| Na<sup>+</sup>-K<sup>+</sup>-ATPase | 0.7 (3) | 0.77 (4) |
| apical P<sub>f</sub>, principal cell | 1 | 0.6 (20) |
| **Inner-medullary collecting duct** |  |  |
| Na<sup>+</sup>-K<sup>+</sup>-ATPase | 0.7 (3) | 0.616 (4) |
| P<sub>f</sub> | 1/5.4 (18,19) | 1/5.4 (18,19) |
| P<sub>urea</sub> | 1/3.4 (18,19) | 1/3.4 (18,19) |
| **Papillary tip** |  |  |
| Interstitial [urea] | 1/2.5 (18) | 1/2.5 (18) |

### Model parameters for hypertension

After a 21-day chronic Ang-2 infusion, Wistar rats do not exhibit a significant change in GFR (7). Consequently, we assume that both single-nephron glomerular filtration rate (SNGFR) and nephron number remain unchanged in both hypertensive male and female rats. Transporter activities in a hypertensive male rat are primarily based on measurements of transporter protein abundance (3). Though the measured increases in abundance of ENaC subunits vary over a wide range, a 21% increase in ENaC activity of hypertensive male rats is assumed to yield Na^+^ excretion consistent with observation (18). Chronic AngII infusion lowers urine osmolality (3), which can be attributed in part to AngII-stimulated thirst and water intake. As such, we lower interstitial urea concentration by a factor of 2.5 in the inner medulla. Interstitial concentrations of other solutes are assumed to remain at baseline values. Further, the increased water intake suppressed arginine vasopressin (AVP) secretion. AVP increases water and urea permeabilities along the inner-medullary collecting duct (19). To account for these changes, we follow the approach in (18) and assume that the tight junction accounts for 30% of total permeabilities in the absence of AVP stimulation. With this assumption, AVP-induced increases in water and urea permeabilities are estimated to be 3.4- and 5.4-fold, respectively. Thus, in the hypertension simulation, inner-medullary collecting duct water and urea permeabilities are reduced by factors of 3.4 and 5.4, respectively.

Transporter activities in a hypertensive female rat are also primarily based on measurements of transporter protein abundance (4,20). Using the reported mean values did not yield sufficiently large increases in urinary volume, sodium excretion, and potassium excretion. As such, most Na^+^ transporter activities were lowered to within one standard deviation of mean values (see Table 1). It is noteworthy that inhibition of SGLT2 in the proximal convoluted tubule is experimentally observed to be coordinated with inhibition of NHE3 (21), so NHE3 activity is assumed to be further decreased. To yield simulation result consistent with experimental excretion (4), we assume in hypertensive female, fold increase of ENaC is significantly higher than in hypertensive male. Changes in inner-medullary collecting duct permeabilities to water and urea, as well as interstitial urea concentration profile, are also adjusted the same way as in the hypertensive male rat model (see above). Lower Na^+^-K^+^-ATPase activity is assumed to reflect the inhibition of Na^+^-K^+^-ATPase in hypertension (22).

### Diuretics simulations

To simulate the effect of loop diuretics effect, we inhibit NKCC2 by 70%. And to simulate the effect of NKCC2 inhibition on the urine concentrating ability of the kidney, we lowered the interstitial concentrations of Na^+^, K^+^, Cl^-^, and urea, as was done in (17) for a normotensive kidney.

The effects of thiazide diuretics and K^+^-sparing diuretics are simulated by 100% inhibition of NCC and ENaC, respectively. Other model parameters are assumed unaffected.

## Results

### Baseline results

We conducted simulations to predict luminal fluid flow, solute concentrations, transcellular and paracellular fluxes, urine output, and solute excretions at steady state, under normotensive and hypertensive conditions. Figure 2 summarizes the predicted segmental fluid, Na^+^, and K^+^ flows, and corresponding segmental transport. In hypertension, Na^+^ transport is reduced along the proximal nephron segments, i.e., the proximal tubule, via the downregulation of NHE3, and the medullary thick ascending limb, via the downregulation of NKCC2 and Na^+^-K^+^-ATPase (Table 1). Consequently, the reabsorption of Na^+^ along these segments decreases, by 7% in male and 9% in female (see Fig. 2A2). In contrast, along nephron segments downstream of the medullary thick ascending limb, which include the cortical thick ascending limb, distal convoluted tubule, connecting tubule, and collecting duct, segmental Na^+^ transport capacity is generally enhanced; activities of NKCC2, NCC, and ENaC are increased, although in female the activities of NHE3 and Na^+^-K^+^-ATPase are assumed lowered (Table 1). With these changes, Na^+^ reabsorption along these segments is predicted to be higher in the hypertensive models compared to normotension, by 38% in male and 35% in female (see Fig. 2A2), indicating a significant downstream shift in Na^+^ transport in hypertension.

**Figure 2.**
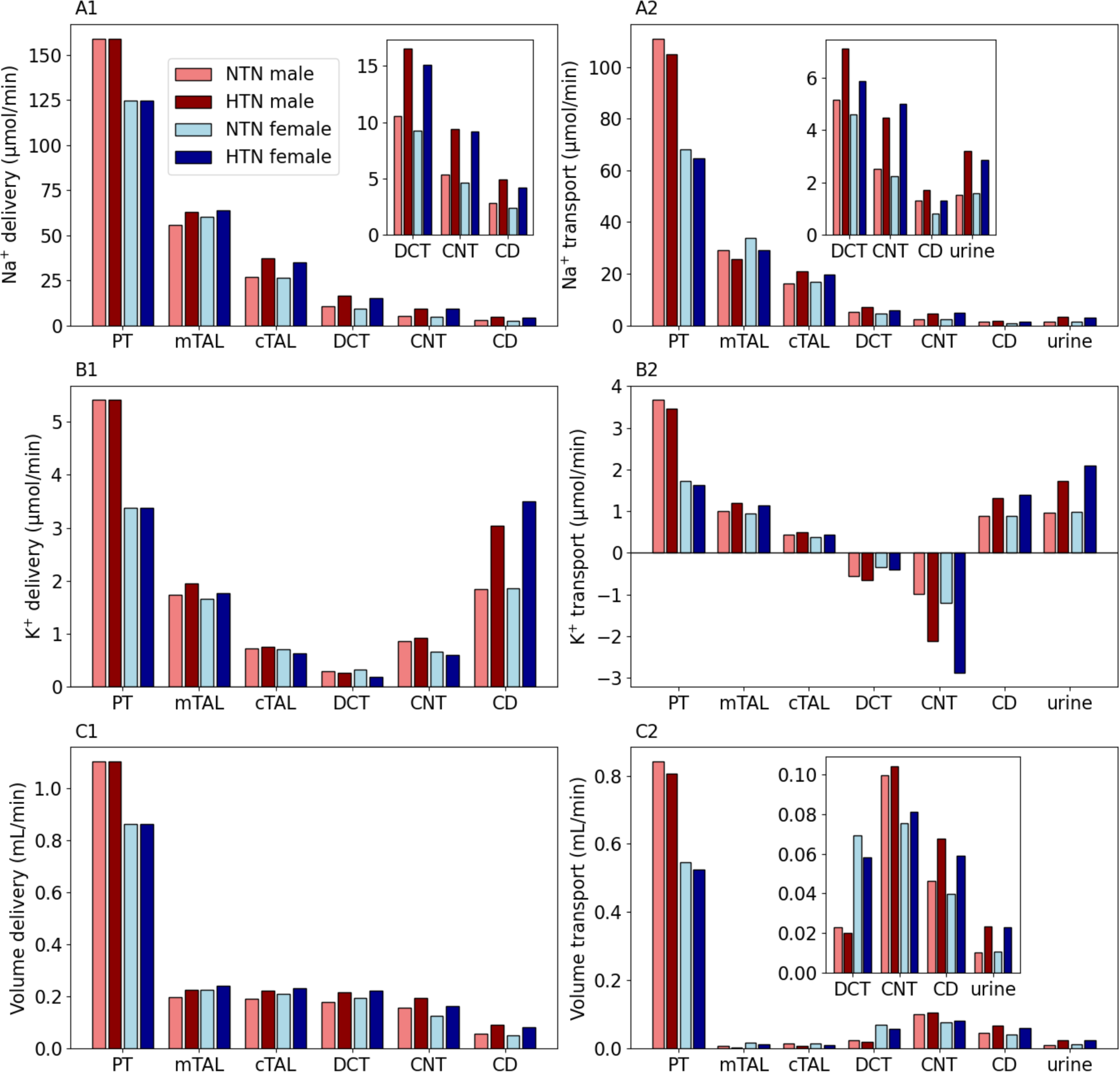
Segmental delivery and transport of Na^+^, K^+^, and water, obtained for the male and female models, under normotensive (NTN) and hypertensive (HTN) conditions. PT, proximal tubule; mTAL, medullary thick ascending limb; cTAL, cortical thick ascending limb; DCT, distal convoluted tubule; CNT, connecting tubule; CD, collecting duct.

Particularly noteworthy is the differential regulation of Na^+^ transport capacity along the thick ascending limb: in hypertension and in both male and female models, NKCC2 activity is reduced along the medullary thick ascending limb but increased along the cortical thick ascending limb (Table 1). As a result, Na^+^ transport is predicted to change in opposing directions along the medullary versus cortical segments (see Fig. 2A2). Those changes mostly cancel out, resulting in a 5% increase in male and a 1% decrease in female from the corresponding normotensive model in overall thick ascending limb Na^+^ reabsorption. These small changes suggest that the differential regulation of Na^+^ transport capacity along the thick ascending limb is not to adjust the segmental Na^+^ reabsorption, but perhaps to adapt to any changes in medullary oxygenation in hypertension (see Discussion). The models predict that an 110% increase in urinary Na^+^ excretion in hypertensive male, compared to normotensive male, and an 80% increase in urinary Na^+^ excretion in hypertensive female above normotensive female.

Water transport is driven primarily by Na^+^ reabsorption; thus, hypertension induces a shift in water transport like that of Na^+^, with water reabsorption significantly decreases along the proximal tubule. and distal convoluted tubule; in contrast, water reabsorption generally increases along the downstream segments. Taken together, in hypertension urine output is predicted to more than double in both sexes (Fig. 2C2).

The reduced water reabsorption in hypertension lowers luminal [K^+^] and thus K^+^ transport along the proximal tubule. AngII-induced proximal tubule NHE3 downregulation results in a significantly lower luminal [NH4^+^] into the downstream segment. While the combined reabsorption of K^+^ and NH4^+^ is lower than in normotension, net K^+^ reabsorption increases in the medullary thick ascending limb.

Along the cortical ascending limb, the upregulated NKCC2 drives higher K^+^ reabsorption. Along the distal convoluted tubule, the lower luminal [K^+^] elevates K^+^ secretion by 16% and 19% above normotension in male and female, respectively. This trend is magnified along the connecting tubule, where K^+^ secretion is higher in hypertension by 115% in male and 142% in female (Fig. 2B2).

Along the collecting duct, K^+^ reabsorption increases by 52% in hypertensive male and 67% in hypertensive female (see Fig. 2B2). Collectively, the models predict 78% and 110% increases in urinary K^+^ excretion in hypertensive male and female, respectively, above normotension.

### NHE3 downregulation in the proximal tubule may be the primary factor contributing to natriuresis and diuresis in hypertension

We assessed the effect of the downregulation of NHE3 in hypertension (Table 1) by conducting simulations in which proximal tubule NHE3 activity is set to normotensive levels. Other hypertension-induced adaptations in the proximal tubule (specifically, the downregulation of Na^+^-K^+^-ATPase and SGLT2 in female) and in other segments are represented. As might be expected, given the essential role of NHE3 in reclaiming filtered Na^+^, returning NHE3 activity to the higher normotensive level substantially reduces Na^+^ excretion to close to normotensive levels in both sexes (see Figs. 3A1, 3A2). The effect on urine output is similar. These results support the downregulation of NHE3 in the proximal tubule as the primary factor contributing to natriuresis and diuresis in hypertension (see Discussion).

**Figure 3.**
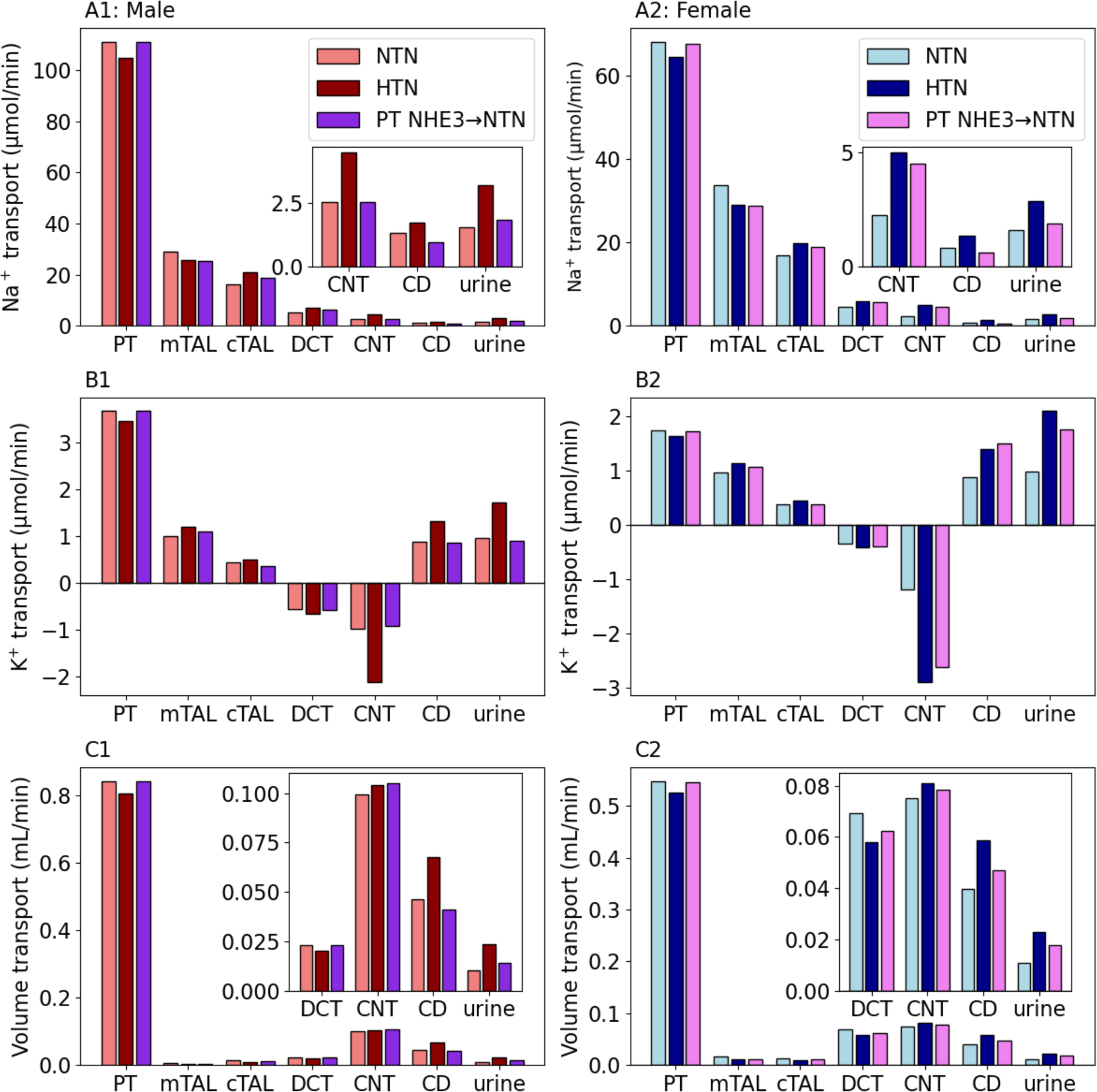
Segmental transport of Na^+^, K^+^, and water in male (A1-C1) and female (A2- C2), obtained for normotension (NTN), hypertension (HTN), and hypertension but with proximal tubule NHE3 activity set to normotensive value (denoted PT NHE3→ NTN). Notations for segments are analogous to Fig. 2.

Eliminating NHE3 downregulation increases proximal reabsorption of Na^+^ and decreases its delivery to distal segment. Consequently, ENaC-mediated Na^+^ is reduced, which in turn attenuates K^+^ secretion. It is noteworthy that the extent to which K^+^ excretion is reduced differs significantly between male and female. In the hypertensive male model without NHE3 downregulation, Na^+^ delivery to the connecting tubule is 43% lower than the baseline hypertensive model, resulting in a substantial 56% decrease in K^+^ secretion along that segment (Fig. 3A2). In the hypertensive female model, Na^+^-K^+^- ATPase and SGLT2 remain downregulated along the proximal tubule. Also, the thick ascending limb accounts for a larger fraction of total Na^+^ transport. Consequently, even with NHE3 activity at the normotensive level, Na^+^ delivery to the connecting tubule is only 24% lower than baseline hypertension (compare to 43% in male), and the resulting reduction in connecting tubule K^+^ secretion is only 10% (compared to 56% in male) as shown in Fig. 3A1. These results suggest that the downregulation of NHE3 in hypertension plays a more important role in hypertension-induced kaliuresis in male than in female.

### Loop diuretics enhance hypertension-induced natriuresis, kaliuresis, and diuresis

We simulate the administration of loop diuretics by inhibiting NKCC2 by 70%. Simulations were conducted using the normotensive and hypertensive models. Results for the male and female models are shown in Fig. 4 and 5, respectively. Results for full NKCC2 inhibition can be found in the Supplemental Materials.

**Figure 4.**
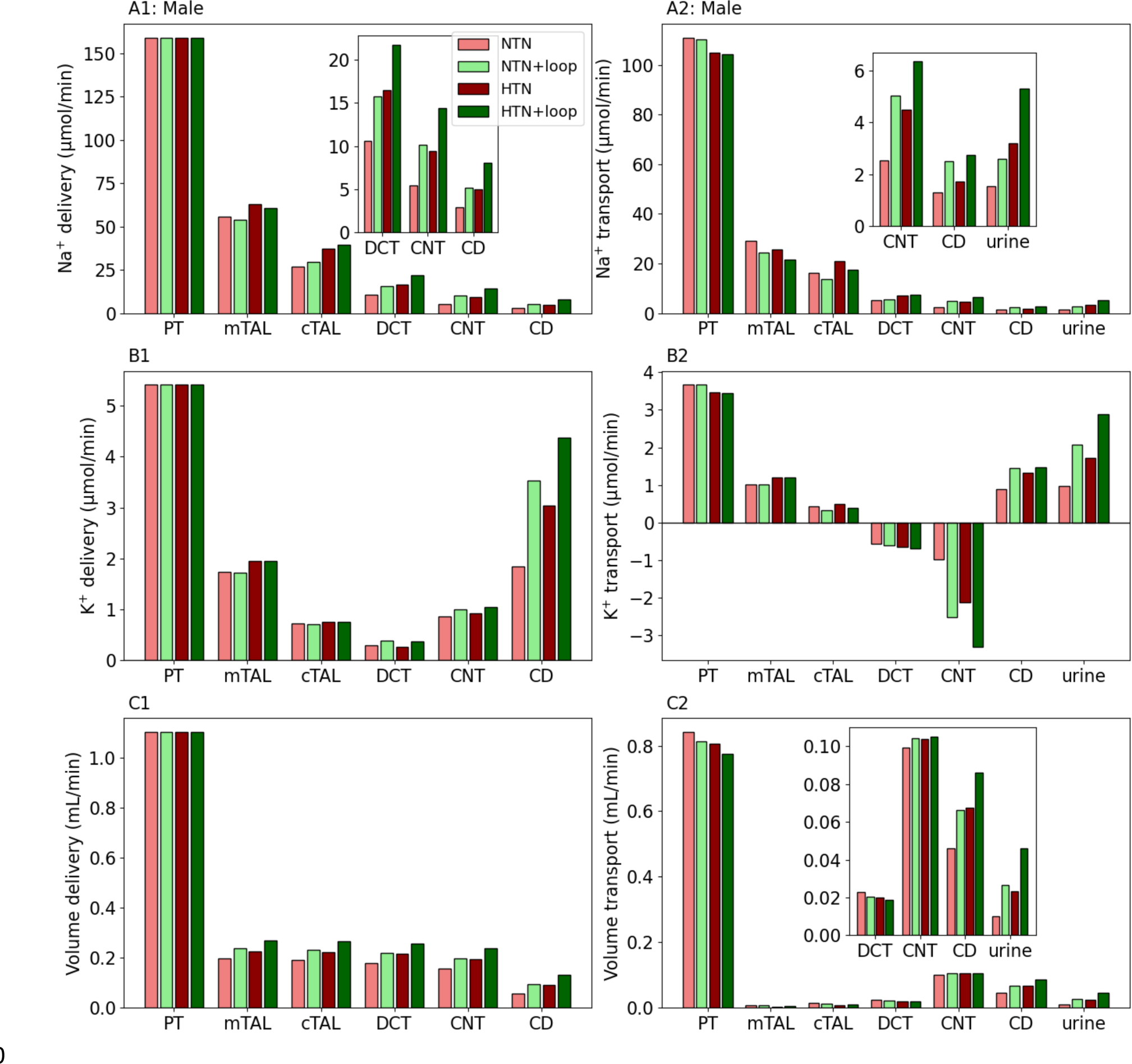
Segmental delivery (A1-C1) and transport (A2-C2) of Na^+^, K^+^, and water in male for normotension (NTN), normotension with 70% NKCC2 inhibition (NTN+loop), hypertension (HTN), hypertension with 70% NKCC2 inhibition (HTN+loop). Notations are analogous to Fig. 2.

**Figure 5.**
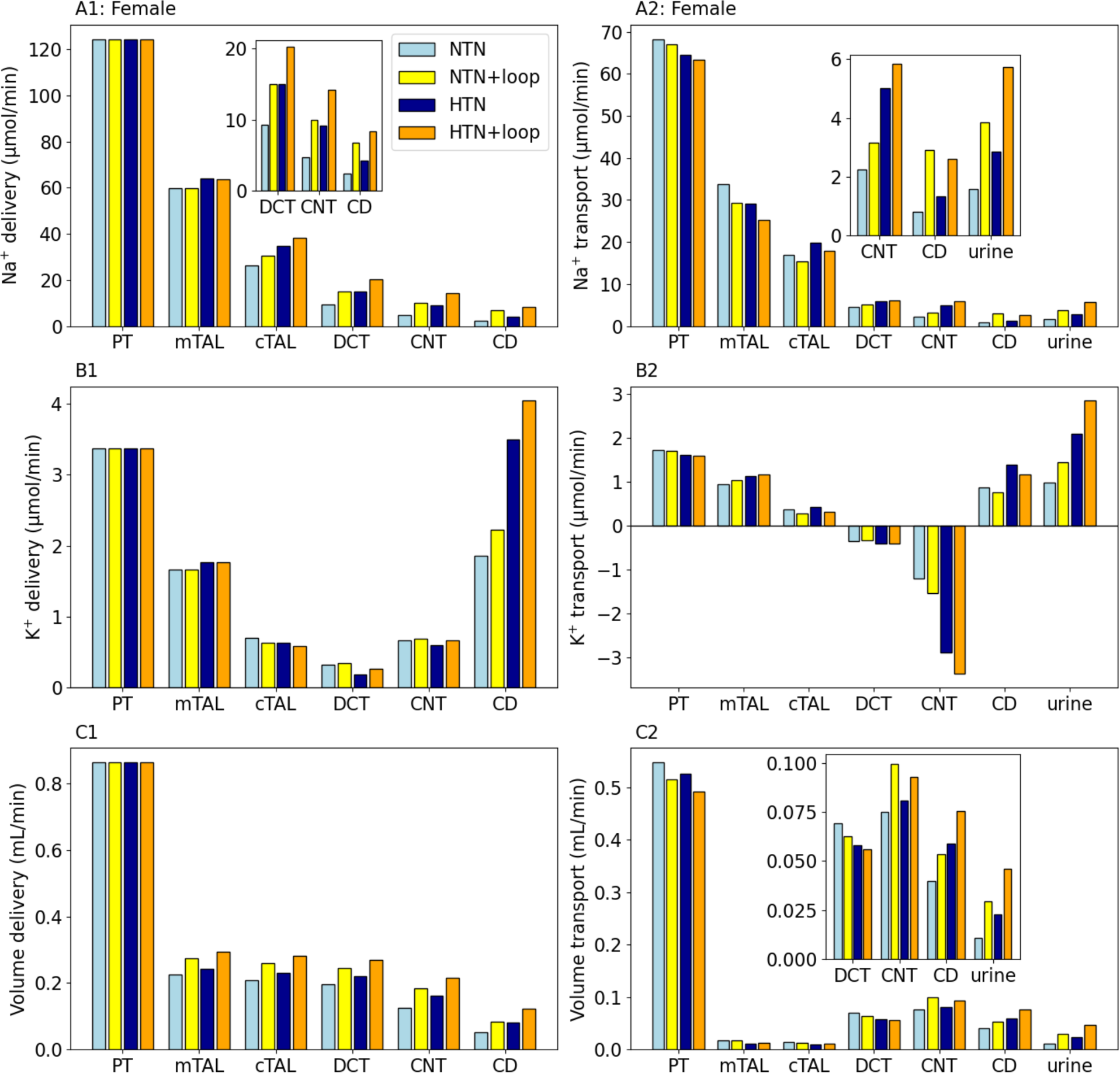
Segmental delivery (A1-C1) and transport (A2-C2) of Na^+^, K^+^, and water in female for normotension (NTN), normotension with 70% NKCC2 inhibition (NTN+loop), hypertension (HTN), hypertension with 70% NKCC2 inhibition (HTN+loop). Notations are analogous to Fig. 2.

With a 70% inhibition of NKCC2, Na^+^ reabsorption along thick ascending limb decreases by 16% in both normotensive and hypertensive male, and by 12% and 11% in normotensive and hypertensive female, from their respective baseline (i.e., without loop diuretics) models. Downstream segments compensate. In particular, Na^+^ reabsorption increases significantly along the connecting tubule (see Figs. 4A2, 5A2). Together, these changes result in substantial increases in Na^+^ excretion (67% and 66% in normotensive and hypertensive male; 140% and 98% in normotensive and hypertensive female), above the respective baseline cases. It is noteworthy that these are *relative changes*. Recall that hypertension induces natriuresis. Thus, while these percentages are smaller in the hypertensive models, especially in female, the administration of loop diuretics is predicted to yield net Na^+^ excretion that is substantially higher in hypertension (with loop diuretics) for both male and female (Figs. 4A2, 5A2), compared to normotension (with loop diuretics).

Partial NKCC2 inhibition is predicted to attenuate K^+^ reabsorption in thick ascending limb, although that reduction is less than that in Na^+^ reabsorption. The higher Na^+^ delivery to the connecting tubule stimulates K^+^ secretion, by 160%, 58%, 26%, and 17% in normotensive male, hypertensive male, normotensive female, and hypertensive female, respectively. Consequently, K^+^ excretion increases by 120% in normotensive male, 65% in hypertensive male, 47% in normotensive female, and 38% in hypertensive female. In both sexes, urinary K^+^ excretion is higher in hypertension than normotension (both with loop diuretics), even the relative increases are smaller in the former (Figs. 4B2, 5B2).

Volume excretions are enhanced in all cases (160% and 98% in normotensive and hypertensive male; 190% and 100% in normotensive and hypertensive female). Because thick ascending limb is essentially water impermeable, the diuretic effect is attributable to the solute concentration increase due to NKCC2 downregulation, which then attenuates water reabsorption downstream.

### Thiazide diuretics induce a stronger natriuretic response in hypertension, especially in females

We simulate the effects of the administration of a thiazide diuretic to the normotensive and hypertensive models, for both males and females (Figs. 6, 7). With NCC fully inhibited, Na^+^ reabsorption along the distal convoluted tubule drops precipitously (Figs. 6A2, 7A2). The reduction in Na^+^ is partially compensated for along the connecting tubule and collecting duct. Nonetheless, the models predict that the inhibition of NCC causes a substantial natriuretic effect, with urinary Na^+^ excretion increases by 52% (40%) and 74% (49%) in the normotensive and hypertensive female (male) models, respectively. The stronger natriuretic effect predicted in the hypertensive models can be attributed, in large part, to the substantially higher Na^+^ flow into the distal segments. Indeed, to handle that larger Na^+^ flow, NCC activity is assumed to essentially double in hypertension in both sexes. When NCC is inhibited by a thiazide diuretic, it is more challenging for the segments downstream of the distal convoluted tubule to compensate in the hypertensive models. A stronger natriuretic response is predicted for the female models. That sex difference may be explained by the relatively limited capacity of the female models’ other Na^+^ transport pathways in the distal convoluted tubule and downstream segments to elevate their Na^+^ transport.

**Figure 6.**
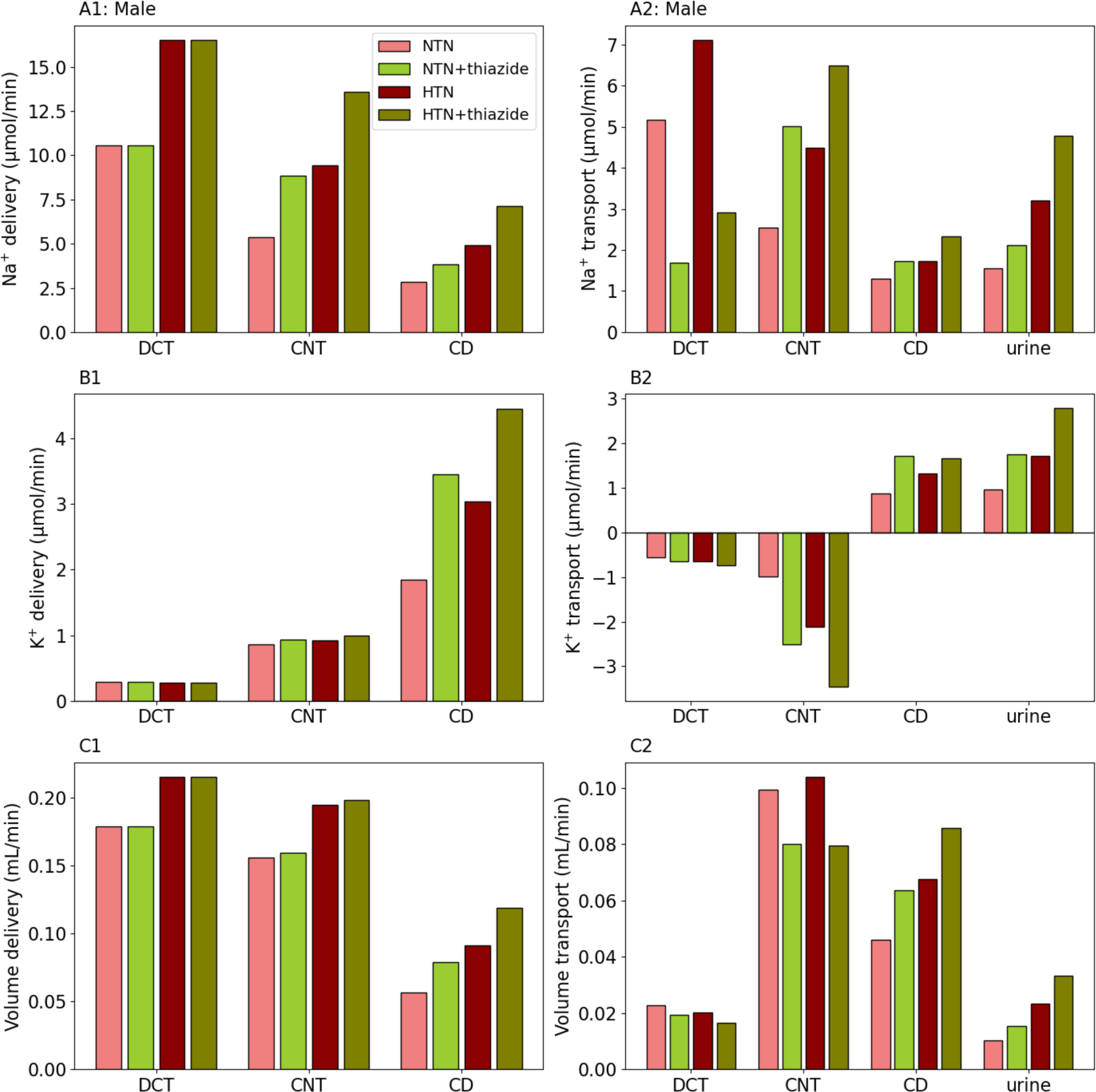
Segmental delivery (A1-C1) and transport (A2-C2) of Na^+^, K^+^, and water in male for normotension (NTN), normotension with 100% NCC inhibition (NTN+thiazide), hypertension (HTN), hypertension with 100% NCC inhibition (HTN+thiazide). Notations are analogous to Fig. 2.

The enhanced ENaC-mediated Na^+^ transport increases K^+^ secretion along the connecting tubule, resulting in kaliuresis (Figs. 6B, 7B). The elimination of NCC- mediated Na^+^ reabsorption along the distal convoluted tubule is followed by a reduction in water reabsorption, although the effect is much smaller (Figs. 6B, 7B). Unlike Na^+^, where the connecting tubule partially makes up for the upstream transport impairment, water reabsorption along the connecting tubule also decreases. Despite some partial compensation along the collecting duct, the models predict substantial diuretic effects (Figs. 6C, 7C).

**Figure 7.**
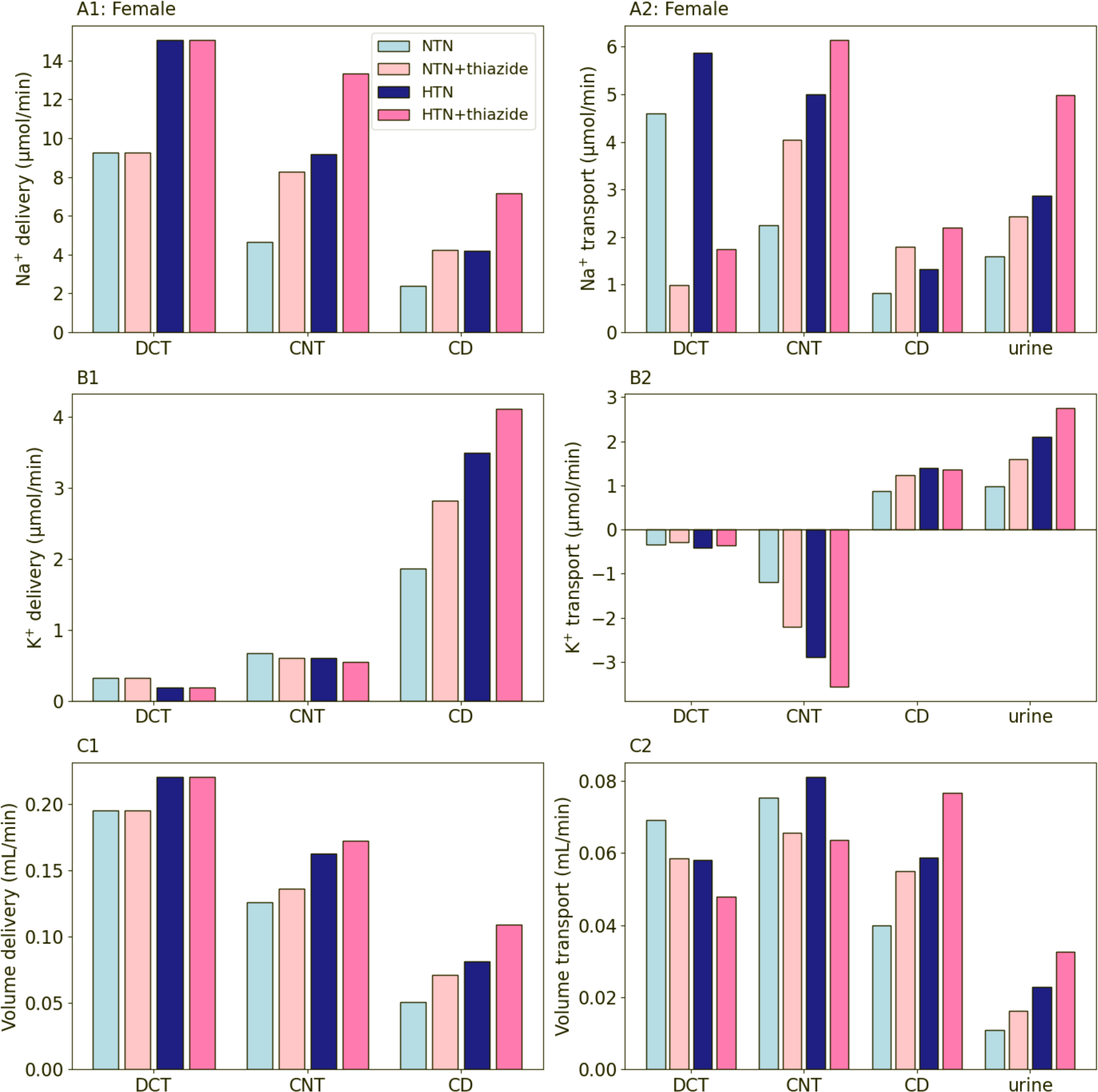
Segmental delivery (A1-C1) and transport (A2-C2) of Na^+^, K^+^, and water in female for normotension (NTN), normotension with 100% NCC inhibition (NTN+thiazide), hypertension (HTN), hypertension with 100% NCC inhibition (HTN+thiazide). Notations are analogous to Fig. 2.

### K^+^-sparing diuretics induce a stronger natriuretic response in hypertension

We then consider the effect of K^+^-sparing diuretics. All models predict that, when ENaC is fully inhibited, Na^+^ transport along the connecting tubule reverses direction, switching from reabsorption in baseline normotension or hypertension to (a small amount of) secretion (see Figs. 8A2, 9A2). Despite some compensation along the collecting duct, the inhibition of ENaC induces a substantial natriuretic effect, with urinary Na^+^ excretion increases by 200% (190%) and 210% (150%) in the normotensive and hypertensive female (male) models, respectively. The stronger natriuretic effect predicted in the hypertensive models is due to the enhanced Na^+^ transport by connecting tubule in hypertension: in both hypertensive male and female models, the connecting tubule reabsorbs about twice as much Na^+^ as the corresponding normotensive models (see Figs. 8A2, 9A2). As such, when that Na^+^ transport capacity is blocked, it results in a larger natriuretic effect in the hypertensive models.

**Figure 8.**
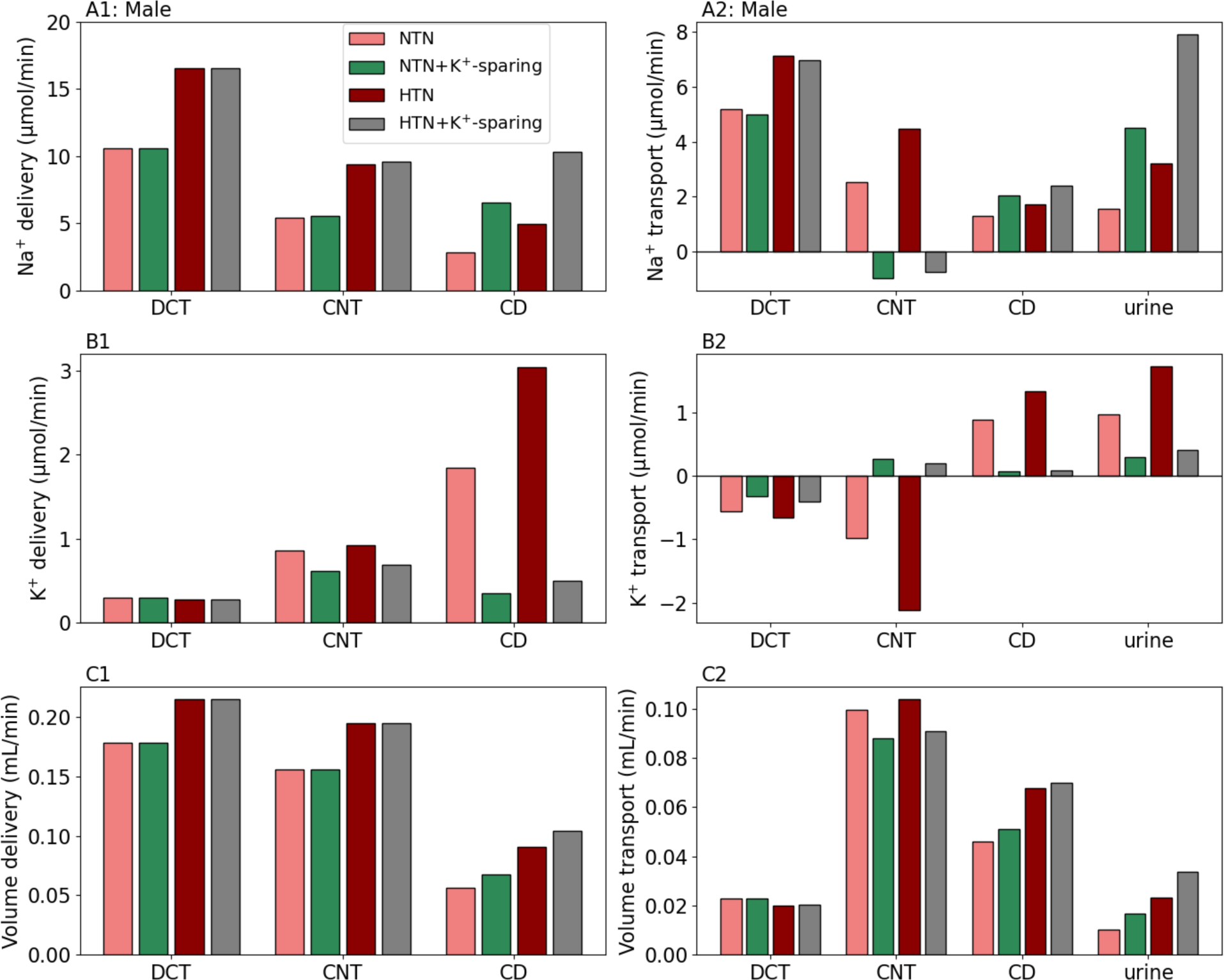
Segmental delivery (A1-C1) and transport (A2-C2) of Na^+^, K^+^, and water in male for normotension (NTN), normotension with 100% ENaC inhibition (NTN+K^+^- sparing), hypertension (HTN), hypertension with 100% ENaC inhibition (HTN+K^+^- sparing). Notations are analogous to Fig. 2.

**Figure 9.**
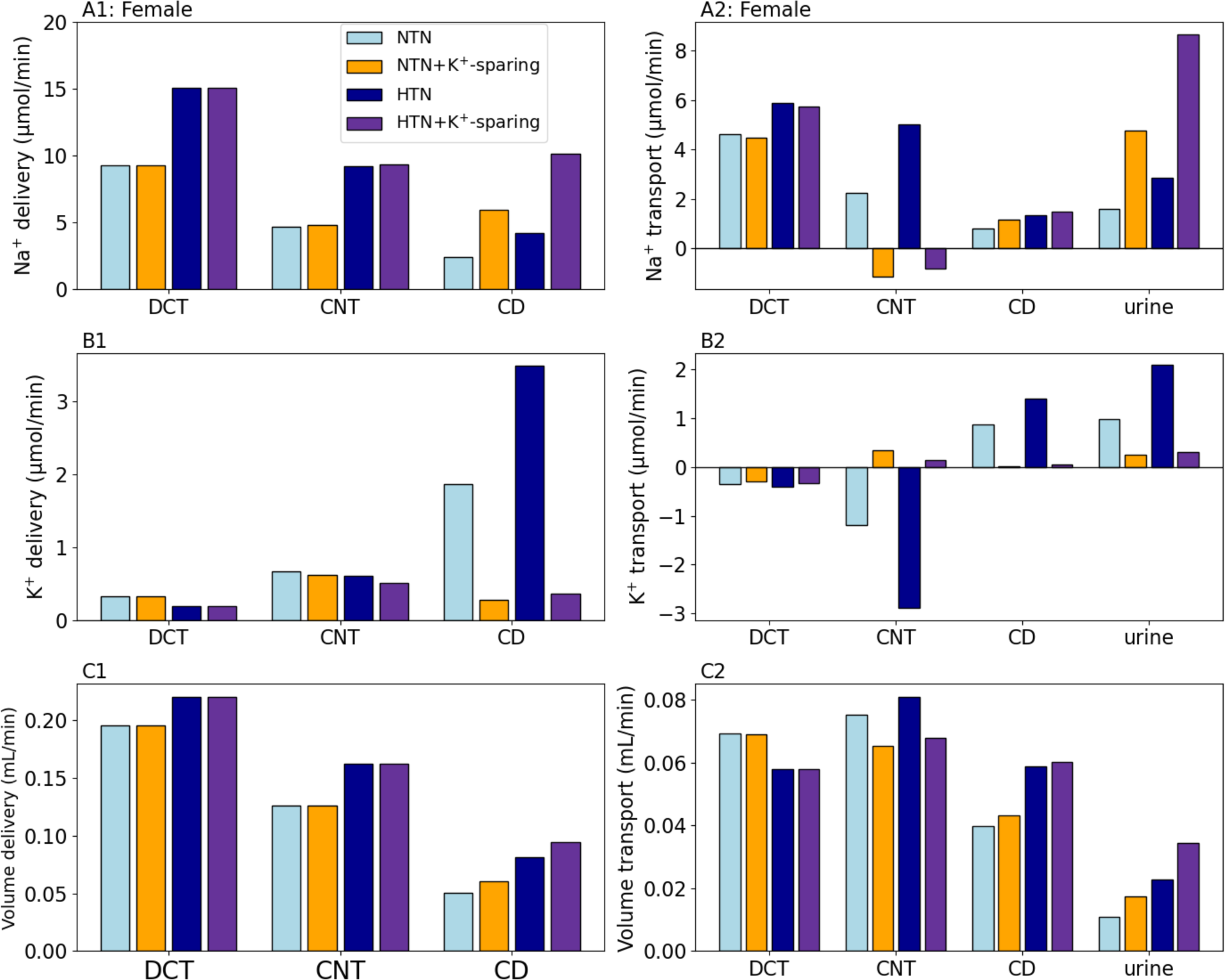
Segmental delivery (A1-C1) and transport (A2-C2) of Na^+^, K^+^, and water in female for normotension (NTN), normotension with 100% ENaC inhibition (NTN+K^+^- sparing), hypertension (HTN), hypertension with 100% ENaC inhibition (HTN+K^+^- sparing). Notations are analogous to Fig. 2.

The removal of ENaC-mediated Na^+^ reabsorption also abolishes K^+^ secretion along the connecting tubule. This results in a substantial reduction of urinary K^+^ excretion. The elimination of Na^+^ reabsorption and K^+^ secretion has opposing effects on water transport. These competing effects result in some reduction in water reabsorption along the connecting tubule (Figs. 8C2, 9C2); eventually, urine output increases by 59% (68%) and 50% (44%) in the normotensive and hypertensive female (male) models, respectively.

## Discussion

In many mammalian and avian species, young males have a higher blood pressure and a higher prevalence of hypertension compared to age-matched females (23,24). This phenomenon is observed not only in humans but also in genetic models of hypertension, including spontaneously hypertensive rats and Dahl salt-sensitive rats, where males tend to experience hypertension at an earlier stage and with more severity than their female counterparts (25). Despite these observations, the precise mechanisms responsible for these differences in blood pressure regulation between males and females remain inadequately elucidated. One of the suggested mechanisms underlying this sexual dimorphism is differences in the function of the RAAS. The RAAS is important in the regulation of body fluid and blood pressure homeostasis, and overactivation of the RAAS contributes to the development of numerous pathophysiological processes, notably, the development and progression of hypertension. There is an expanding literature regarding sex differences in the expression of RAAS components (26–29), as well as in functional responses to RAAS activation to both acute and chronic AngII infusion (30).

A widely utilized experimental model of hypertension is the low-dose infusion of AngII via osmotic minipump at 400 ng/kg/min (2,31–33). The kidney’s response to the chronic infusion of AngII, a Na^+^-retaining hormone, is biphasic. In male rodents, 3-day low-dose infusion of AngII stimulates the activation of sodium transporters all along the nephron (34). With continued AngII infusion, pressure natriuresis takes effect to reinstate fluid and electrolyte homeostasis, whereby proximal and medullary NHE3 and medullary NKCC2 pool sizes decrease, while cortical NKCC2, NKCC2p, NCC, NCC-p, and cleaved ENaC subunits remain activated (3,4). Notably, the shift of Na^+^ transport capacity to distal segments in AngII-induced hypertensive male rodents renders their transporter more similar to that of normotensive females, allowing for better fine-tuning of Na^+^ transport. AngII infusion in female C57BL/6J mice amplified the baseline profile by decreasing the pool size of NHE3 and increasing the pool sizes of NCC and ENaC (4).

Model simulations indicate that this beneficial pattern, with the lower proximal and medullary thick ascending limb Na^+^ transporter abundance along with enhanced distal and collecting duct transport capacity, facilitates pressure natriuresis by significantly suppressing proximal and medullary thick ascending limb Na^+^ transport. Model simulations predict that fractional Na^+^ transport along proximal tubule and medullary thick ascending limb in male and female rat nephrons decreases by approximately 10%. Most of that reduction can be attributed to the inhibition of NHE3 along the proximal tubule. Without that adaptation, the natriuretic effect would be much attenuated, with urinary Na^+^ excretion decreased by about 1/3 from the baseline hypertensive models (Figs. 3A1, 3A2).

Oxygen tension is lower in the renal medulla than in renal cortex (35). AngII-induced vasoconstriction can lead to pathogenesis of pulmonary hypertension (36), which may result in hypoxemia (37). Because tubular Na^+^ reabsorption is a major contributor of oxygen consumption in kidney (38,39), a partial redistribution of the workload required for Na^+^ reabsorption from the renal medulla to the cortex may serve to avoid severe medullary hypoxia. Upregulation of ENaC and NCC shifts Na^+^ reabsorption load from the collecting duct segment in a region with the lowest oxygen tension (inner medulla) to the outer medullary collecting duct (40).

Increased urinary Na^+^ excretions are observed in both humans with essential hypertension (41,42) and rodents with spontaneous (43) or AngII-induced hypertension (4,34). Model simulations predict that female rats with AngII-induced hypertension have lower increase in urinary Na^+^ excretions than that in male (Fig. 2A2), consistent with experimental studies showing that female rodents have lower mean arterial pressure and relative increase in Na^+^ excretions than males, following AngII infusion (3,4).

Indeed, females appear to be more resistant to AngII-induced hypertension, a sex difference that may be attributed to higher bioavailability of nitric oxide in females (15,44) and a less reactive RAAS (45).

In separate studies, Edwards et al. applied computational models of epithelial transport in nephrons to study the impact of 14-day AngII infusion on the handling of Na^+^ and K^+^ in male (18) and female (20) rats. Their simulations predict that in both sexes AngII infusion stimulates distal Na^+^ reabsorption and K^+^ secretion, and that a substantial downregulation of proximal tubule NHE3 is needed to reestablish Na^+^ balance at 2 weeks. As discussed above, a similar downstream shift in Na^+^ reabsorptive and K^+^ secretive loads is seen in our simulation results. A comparison between the predicted urinary excretion (Fig. 3) indicates that the contribution of hypertension-induced downregulation of PT NHE3 activity to downstream shift of Na^+^ transport is stronger in the hypertensive male model than female. When PT NHE3 activity was returned to the (higher) normotensive value, the male model predicted substantially lower urine output and Na^+^ and K^+^ excretion rates that are close to normotensive excretions (Fig. 3). In contrast, K^+^ excretion in particular is notably higher than the normotensive excretion and closer to the hypertensive value in the female model (Figs. 3B1, 3B2). That sex difference can be attributed to the hypertension-induced transporter alterations in the PCT. In males, that is limited to a reduction in NHE3 activity, whereas in females, Na^+^-K^+^-ATPase and SGLT2 activities are also downregulated. Our simulations of the administration of diuretics on nephron transport in a hypertensive kidney indicate that the natriuretic and diuretic effects are generally stronger in females, consistent with experimental findings (46–48); more below.

### Loop diuretics

Expressed on the apical membrane of the thick ascending limb and macula densa cells, the bumetanide-sensitive NKCC2 facilitates the primary apical pathway for the reabsorption of Na^+^ and Cl^-^ along the thick ascending limb. NKCC2 can be inhibited by loop diuretics such as furosemide, a widely used, potent natriuretic drug that can be administered to mitigate extracellular fluid volume expansion associated with heart and kidney diseases.

Micropuncture studies by Hropot et al. (49) on male Sprague-Dawley rats reported an approximately four-fold increase in early distal Na^+^ delivery after the administration of furosemide. Reyes reported that administration of furosemide to healthy adults yielded dose-dependent natriuresis (50). In humans, dysfunction of NKCC2 in humans is linked to type 1 Bartter syndrome (51), a nephropathy that involves severe salt-wasting and the inability to generate concentrated urine. A mouse model of Bartter’s syndrome exhibit polyuria (52). Consistent with these observations, model simulations suggest that a partial (70%) inhibition of NKCC2 in normotension leads a substantial increase in early distal delivery of Na^+^ (Figs. 4A1, 5A1), resulting in approximately 0.67- to 1.4-fold increase in urinary Na^+^ excretion. A completely inhibition of NKCC2 in normotension yields 8.6- to 18-fold increase in Na^+^ excretion, coupled with a rise of approximately 2.0- to 4.3-fold in early distal Na^+^ flow (see Supplemental Materials Fig. S1A1, S2A1). Alterations of segmental Na^+^, K^+^, and water transport are predicted to be similar in hypertension, with segmental transport shifted downstream, and a magnification of the natriuresis, diuresis, and kaliuresis induced in hypertension.

Despite some promising observations in treatment of fluid overload, uncertainties remain regarding the intrarenal and systemic actions of furosemide. NKCC2 inhibition affects loop of Henle transport and macula densa sensing of Cl^-^, which has implications on the function of the tubuloglomerular feedback mechanism and the regulation of renin release. Indeed, how NKCC2 expression on the macula densa cell membrane is altered in AngII-induced hypertension is not known. Due to the incomplete knowledge of these interacting factors, the GFR responses to furosemide in healthy subjects appear to be variable (53). Further complicating its use is the erratic absorption of furosemide, with a bioavailability ranging from 12% to 112% (54).

### Thiazide diuretics

NCC plays a crucial role in kidney function, specifically electrolyte and fluid homeostasis by mediating Na^+^ and Cl^-^ reabsorption in the initial segment of the distal convoluted tubule. Mutations in the NCC result in Gitelman syndrome, a salt-wasting disorder (55). Given its major role in the regulation of salt, fluid, blood pressure, NCC is a target for a class of antihypertensive medications, thiazide diuretics (56). Thiazide diuretics, an FDA-approved drug class, target NCC and impede the reabsorption of 3% to 5% of luminal Na^+^ in the distal convoluted tubule of the nephron (57). This action facilitates natriuresis and diuresis. Commonly utilized thiazide diuretics include hydrochlorothiazide (HCTZ), chlorthalidone, and indapamide (58).

Inhibition of NCC results in substantially increased delivery of Na^+^ to the distal segment of the distal convoluted tubule and collecting tubule (Figs. 6A1, 7A1). The higher Na^+^ flow enhances Na^+^ reabsorption in the principal cells via ENaC on the apical membrane and the aldosterone-sensitive Na^+^-K^+^-ATPase pump on the basolateral membrane. The enhanced Na^+^ flux generates an electrical potential favorable for K^+^ secretion into the lumen. Model simulations predicted considerable increases in urinary K^+^ excretion (Figs. 6B2, 7B2). Indeed, a drop in serum K^+^ is a well-known effect of thiazide diuretics (59). It is estimated that ≤50% of patients receiving thiazide-type diuretics develop hypokalemia (defined as a serum K^+^ <3.5 mmol/L) (60).

### K^+^-sparing diuretics

The final adjustment of Na^+^ reabsorption occurs along the distal segments, which include terminal segment of the distal convoluted tubule, the connecting tubule, and the collecting duct. Along these segments, Na^+^ reabsorption is mediated by the amiloride-sensitive ENaC (61), located on the apical membrane. Loss-of-function mutations in ENaC occurs in pseudohypoaldosteronism type 1, the dominant form of which is very severe (62). Gain-of-function mutations in ENaC leads to Liddle syndrome, which is characterized by hypertension and hypokalemic alkalosis associated with hypoaldosteronemia (63).

Under most conditions, loop diuretics are K^+^-wasting, and patients may require K^+^ supplementation. Remarkably, Wang et al. (64) demonstrate that in mice fed a low- Na^+^, high-K^+^ diet, loop diuretics are K^+^-sparing. Normally, K^+^ is absorbed along the thick ascending limb. However, when the mice are fed a low-Na^+^, high-K^+^ diet, their thick ascending limb of these mice mediate significant K^+^ secretion. That switch is driven by the elevated activities of NKCC2 and Renal Outer-Medullary K^+^ channels (ROMK), which provide an electrochemical gradient that favors K^+^ secretion, transforming loop diuretics into K^+^-sparing agents. It is interesting to note that the low-Na^+^, high-K^+^ diet is reminiscent of the diet consumed by earlier humans.

### Model limitations and future extensions

The goal of this study is to develop computational tools for conducting *in silico* experiments to advance our understanding of the kidney’s role in pathogenesis and treatments of hypertension. As we apply the models to conduct simulations and analyze data, it is important to understand and acknowledge their limitations. These limitations include the scarcity of direct measurements of transport activities, and the limited availability of experimental data at the cellular level, including details of intracellular signaling pathways, especially in female species (65). Additionally, the current models represent a superficial nephron, which account for two-thirds of the total nephron population of a rat kidney (66). Focus on the superficial nephron allows us to better understand the impact of hypertension-induced alternations in individual transporters. The rest of the nephrons are juxtamedullary one, which can be incorporated in a more comprehensive computational model of kidney function to provide more accurate predictions of whole-kidney function (e.g., Refs. (67,68)).

The present models are formulated for AngII-induced hypertension, which accounts for a disproportionally large fraction (nearly half) of published hypertension animal research studies (31). Other forms of hypertension, e.g., genetic animal models of hypertension, mineralocortical-salt hypertension, L-NAME-induced hypertension, etc. likely have different effects on kidney function. However, sex-specific renal transporter measurements are mostly unavailable in these animal models. Furthermore, people with hypertension often suffer from other diseases such as diabetes, which also impact kidney function (69,70). Understanding kidney function in pregnancy (71,72) and hypertension may lead to better treatment options for gestational hypertension and preeclampsia.

## Notes

### Competing Interest Statement

The authors have declared no competing interest.

