## Supplement Text for "Modeling Sex Differences in the Effects of Diuretics in Renal Epithelial Transport during Angiotensin II-induced Hypertension"

### Supplement Materials

*100% NKCC2 inhibition (high dosage of loop diuretics) induces severer side effects in hypertension than in normotension.*

High dosage of loop diuretics may lead to side effects especially dehydration (1). Results for a dosage that inhibit 70% of NKCC2 are reported in the main text. Here we consider a dosage is sufficiently high to cause 100% inhibition of NKCC2. Simulations are conducted for male and female under both normotension and hypertension. Results are summarized in Figs. S1-S2.

In all cases, 100% NKCC2 inhibition is predicted to reduce $Na^+$ reabsorption to below 20% of baseline case (i.e., without diuretics) along the entire thick ascending limb. $K^+$ transport reverses direction, switching from reabsorption to secretion. The natriuresis, kaliuresis, and diuresis responses are higher in the male and female hypertensive models compared to the respective normotensive models. In the hypertensive male model, the fractional excretions of $Na^+$, $K^+$, and water are predicted to be 17%, 120%, and 20%, respectively; for hypertensive females, the corresponding fraction excretion rates are 28%, 160%, and 29%. This may imply high risks of hyponatremia, dehydration, and hypokalemia in hypertensive females.

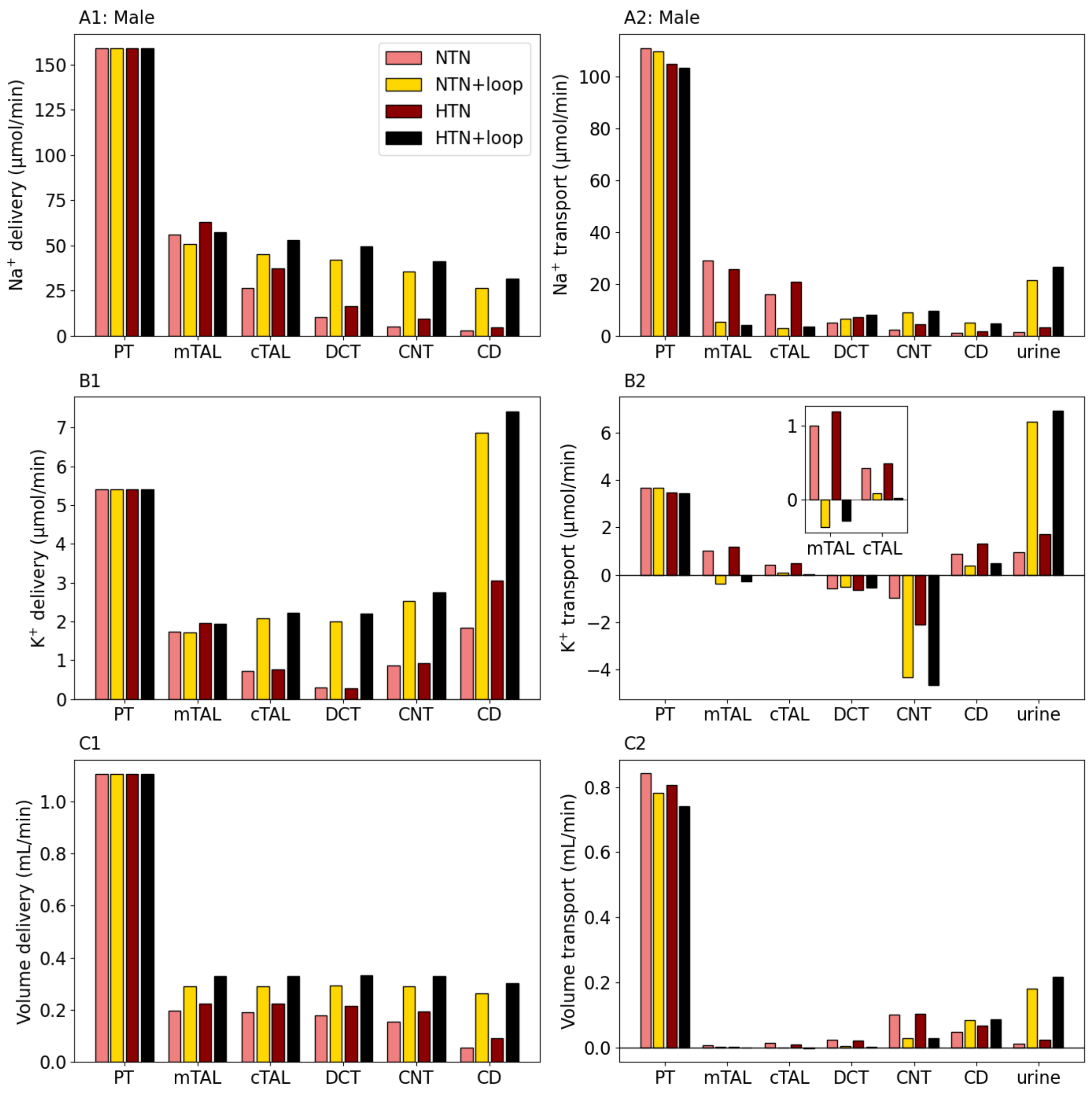


**Figure S1.** Segmental delivery (A1-C1) and transport (A2-C2) of $Na^+$, $K^+$, and water in male for normotension (NTN), normotension with 100% NKCC2 inhibition (NTN+loop), hypertension (HTN), hypertension with 100% NKCC2 inhibition (HTN+loop). PT, proximal tubule; mTAL, medullary thick ascending limb; cTAL, cortical thick ascending limb; DCT, distal convoluted tubule; CNT, connecting tubule; CD, collecting duct; Urine: urinary excretion.

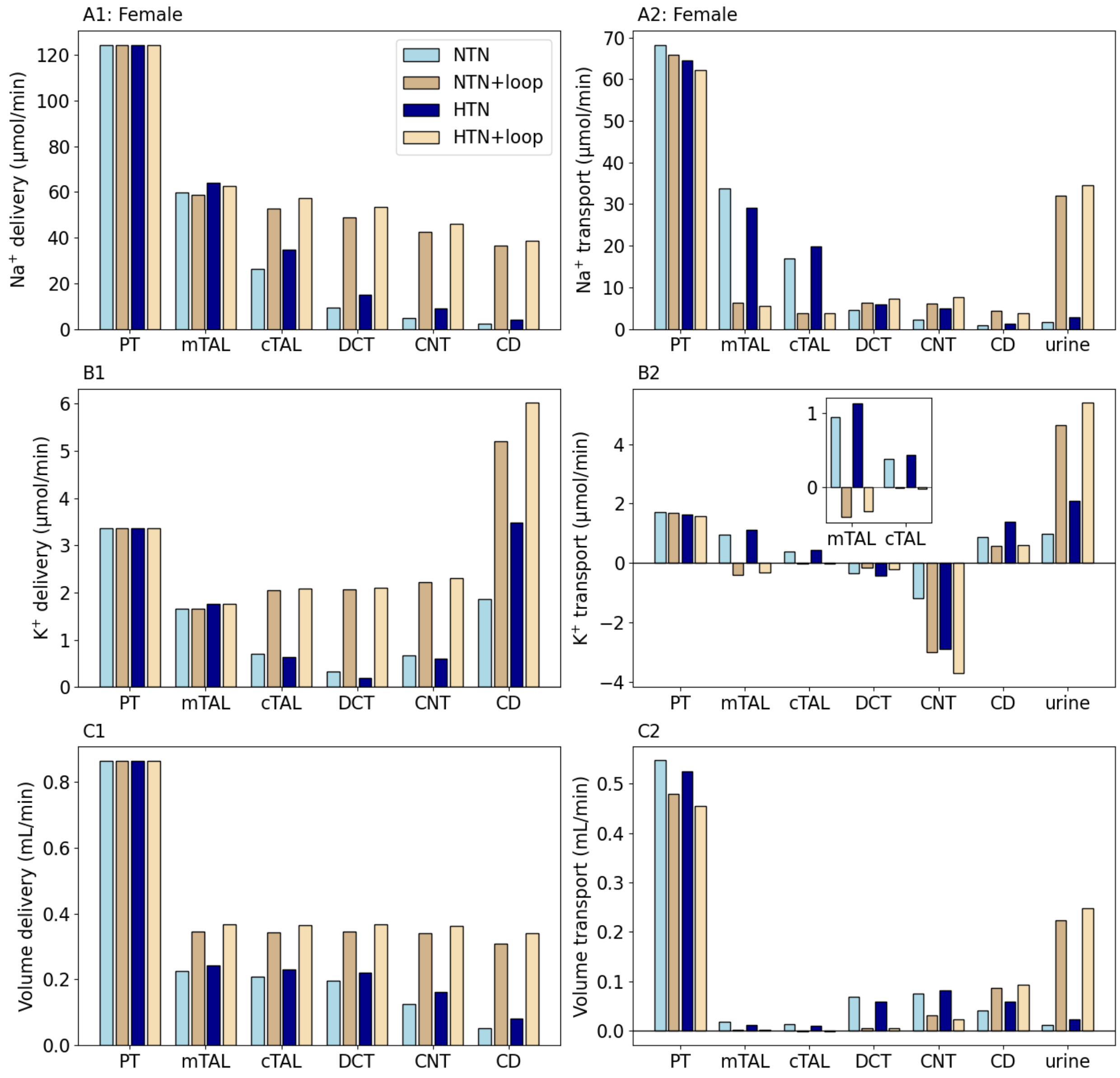


**Figure S2.** Segmental delivery (A1-C1) and transport (A2-C2) of $Na^+$, $K^+$, and water in female for normotension (NTN), normotension with 100% NKCC2 inhibition (NTN+loop), hypertension (HTN), hypertension with 100% NKCC2 inhibition (HTN+loop). Notations are analogous to Fig. S1.
